# Lipid nanoparticles delivering constitutively active STING mRNA as a novel anti-cancer therapeutic approach

**DOI:** 10.1101/2022.01.08.475499

**Authors:** Wei Liu, Mohamad-Gabriel Alameh, June F. Yang, Jonathan R. Xu, Paulo JC Lin, Ying K Tam, Drew Weissman, Jianxin You

## Abstract

Treating immunosuppressive tumors represents a major challenge in cancer therapies. Activation of STING signaling has shown remarkable potential to invigorate the immunologically ‘cold’ tumor microenvironment (TME). However, we and others have shown that STING is silenced in many cancers, including pancreatic ductal adenocarcinoma (PDAC) and Merkel cell carcinoma (MCC), both of which are associated with an immune-dampened TME. In this study, we applied mRNA lipid nanoparticles (LNP) to deliver a permanently active gain-of-function STING^R284S^ mutant into PDAC and MCC cells. Expression of STING^R284S^ induces cytokines and chemokines crucial for promoting intratumoral infiltration of CD8^+^ T cells and, importantly, also leads to robust cancer cell death while avoiding T cell entry and toxicity. Our studies demonstrated that mRNA-LNP delivery of STING^R284S^ could be explored as a novel therapeutic tool to reactivate antitumor response in an array of STING-deficient cancers while overcoming the toxicity and limitations of conventional STING agonists.

## Introduction

Tumor immune suppression represents a major obstacle in achieving effective cancer immunotherapy. This is a clinical challenge present in many human malignancies, including pancreatic cancer. Pancreatic cancer causes the death of around 430,000 patients per year and persists as one of the deadliest malignancies in the world [1–4]. Few effective treatments are available for patients with advanced pancreatic cancer [5]. Nearly 98% of pancreatic cancer patients are also resistant to PD-1/PD-L1 immune checkpoint blockade therapies [6–9]. Thus, there is a significant unmet need for developing more effective therapies targeting this highly lethal cancer.

Pancreatic cancer often establishes a highly immunosuppressive tumor microenvironment (TME), which hinders retaliation by the host immune system and resists immunotherapies [2, 10]. Therefore, cases of this cancer are traditionally classified as non-immunogenic “cold” tumors [2,10,11]. Typically, tumor-infiltrating CD8^+^ cytotoxic T cells are strongly associated with patient survival. However, the majority of pancreatic cancers lack successful infiltration of effective CD8^+^ T cells in the TME [11–13]. Poor intratumoral T cell infiltration and activation present a major hurdle for developing effective immunotherapies, revealing the need for novel therapeutic strategies.

In our previous studies, we discovered that repression of Stimulator of interferon genes (STING) is a key factor underpinning the immunologically “cold” TME of Merkel cell carcinoma (MCC) [14], which is another highly aggressive cancer with over 30% of patients showing metastatic disease at first presentation [15, 16]. STING is a key regulator of innate immune signaling and antitumor responses [17–20]. The canonical role of the STING signaling pathway is to sense cytoplasmic double-stranded DNA (dsDNA) including host cytoplasmatic chromatin, mitochondrial DNA, and foreign dsDNA such as viral dsDNA. These DNA molecules are recognized by cyclic GMP-AMP synthase (cGAS), which in turn synthesizes 2′3′-cGAMP that can bind to and activate STING. After stimulation by pathogen- or damage-associated molecular patterns (PAMPs or DAMPs), STING activates the transcription of type I and III interferons (IFNs) and other pro-inflammatory cytokines to initiate the innate immune response [21–24]. Cancer cells often maintain abundant damaged DNA, which can also stimulate STING-dependent induction of IFNs and several other anti-tumor cytokines/chemokines including CXCL10 and CCL5 [17,20–23]. Among the molecules activated by STING signaling, IFNs can stimulate the generation of anti-tumor T cells, T-cell infiltration, and the direct killing of cancer cells [25–27], whereas CXCL10 and CCL5 are important for recruiting tumor-reactive effector T cells [17-20,28-30]. Therefore, activation of the STING signaling pathway has demonstrated great promise to trigger a switch in the TME of tumors from an immune suppressed ‘cold’ environment to an immune activated ‘hot’ environment [14,17–20,29–36].

We recently discovered that STING is silenced in MCC and that reactivating STING stimulates antitumor inflammatory cytokine/chemokine production, cytotoxic T cell infiltration and activation, and eradication of MCC cells [14]. Our studies in MCC provide proof-of-principle data to support the hypothesis that targeted reactivation of STING can bolster antitumor cytotoxicity and invigorate the immune-dampened TME in STING-silenced and immunologically ‘cold’ tumors. We also found that STING is silenced or downregulated in a number of other types of cancers, such as pancreatic ductal adenocarcinoma (PDAC) [14]. In this study, we set out to develop new strategies to reactivate STING signaling in order to bolster antitumor immunity and enforce tumor immunogenicity in STING-silenced PDAC cancer.

In light of the recent findings on the antitumor functions of the cGAS-STING pathway, several STING agonists have been developed to stimulate anti-tumor immune responses, including the activation of CD8^+^ T-cells and natural killer cells, and to induce tumor regression in multiple mouse tumor models [19,32,36–38]. However, clinical trials of these STING agonists did not show beneficial results [39, 40]. The contradicting outcomes between mouse tumor models and human clinical trials might be associated with the distinct levels of *STING* gene expression in these different tumors. While STING is highly expressed in many mouse tumor cell lines, such as CT26 [41] and B16-F10 [42], it is silenced in several human cancers as demonstrated in studies from our group and others [14,20,43]. It has also been shown that B16-F10 *STING* knockout cells were more resistant to immunotherapy [42]. Thus, although STING has been shown to be a promising target for cancer therapies, STING agonists might be of little benefit in STING-deficient tumors.

To overcome the limitations of traditional STING agonists, which do not work in STING-silenced cancers [36,44,45], we explored the idea of introducing naturally occurring constitutively active gain-of-function STING mutants [46, 47] into STING-silenced immunologically “cold” PDAC to reactivate antitumor immunity. STING gain-of-function mutations have emerged in multiple systemic autoinflammatory diseases, including STING-associated vasculopathy with onset in infancy, systemic lupus erythematosus-like syndromes, and familial chilblain lupus diseases [46–61]. These mutations support constitutively hyperactive STING activity, which induces an excessive IFN response that attracts and amasses proinflammatory cells to cause autoimmune disease symptoms [46-51,54,56-60]. We therefore hypothesized that these gain-of-function STING (‘hot’ STING) mutants could be leveraged to activate the STING signaling pathway for treating STING-deficient cancers. In this study, we explored the idea of harnessing these permanently active ‘hot’ STING mutants to ‘heat up’ STING-deficient immunologically ‘cold’ cancers.

We first discovered that the expression of the STING^R284S^ mutant in PDAC cells robustly activates the STING signaling pathway. To further develop a ‘hot’ STING cancer therapy, we generated lipid nanoparticles (LNP) to deliver STING^R284S^ mRNA into cells. We observed that the LNP-delivered STING^R284S^ mRNA could vigorously reactivate anti-tumor cytokine production and induce cancer cell death in STING-silenced PDAC and MCC cells. Moreover, because T cells are intrinsically resistant to exogenous mRNA delivery by LNP, STING^R284S^ mRNA-LNP do not introduce T cell cytotoxicity, which could normally be induced by traditional STING agonists. Our results suggest that STING^R284S^ mRNA-LNP can overcome the toxicity and limitations of conventional STING agonists and therefore could be exploited as a new therapeutic approach for treating an array of STING-deficient cancers that are refractory to current therapies.

## Results

### STING is downregulated in some PDAC lesions

We recently discovered that STING expression is absent in MCC and several other cancer cells, including a number of PDAC cell lines [14]. Following up on that study, we analyzed the STING protein levels in several PDAC cell lines and patient lesions (Fig. 1). From the cell line analysis, we found that STING protein is scarce in AsPC-1, PANC-1, and Capan-1 cells, and virtually undetectable in MIA PaCa-2, as compared with primary human dermal fibroblasts (HDFs). In contrast, the levels of cyclic GMP-AMP synthase (cGAS), the upstream activator of STING, are clearly detected in all the tested cell lines (Fig.1A). To confirm these observations, we co-stained STING protein and the PDAC marker CK19 [62] to examine the STING protein level in PDAC tumor lesions. STING was nearly untraceable in three out of the seven lesions, including those from patients #1780, #4476, and #4021. An interesting observation was made for the lesions isolated from patients #T5_1589 and #3917: while STING signal was detected in CK19^-^ cells, it was found to be specifically silenced in CK19^+^ cells (Fig.1B). The rest of the PDAC lesions, from patient #3791 and patient #1832, indicate normal STING protein level (Fig.1B). These results demonstrate that STING expression could be silenced or repressed in certain PDACs and there appeared to be a pattern of tumor cell-specific repression in some PDAC lesions. Our finding suggests that STING downregulation may contribute to the immunologically ‘cold’ TME in some PDACs.

**Figure 1.**
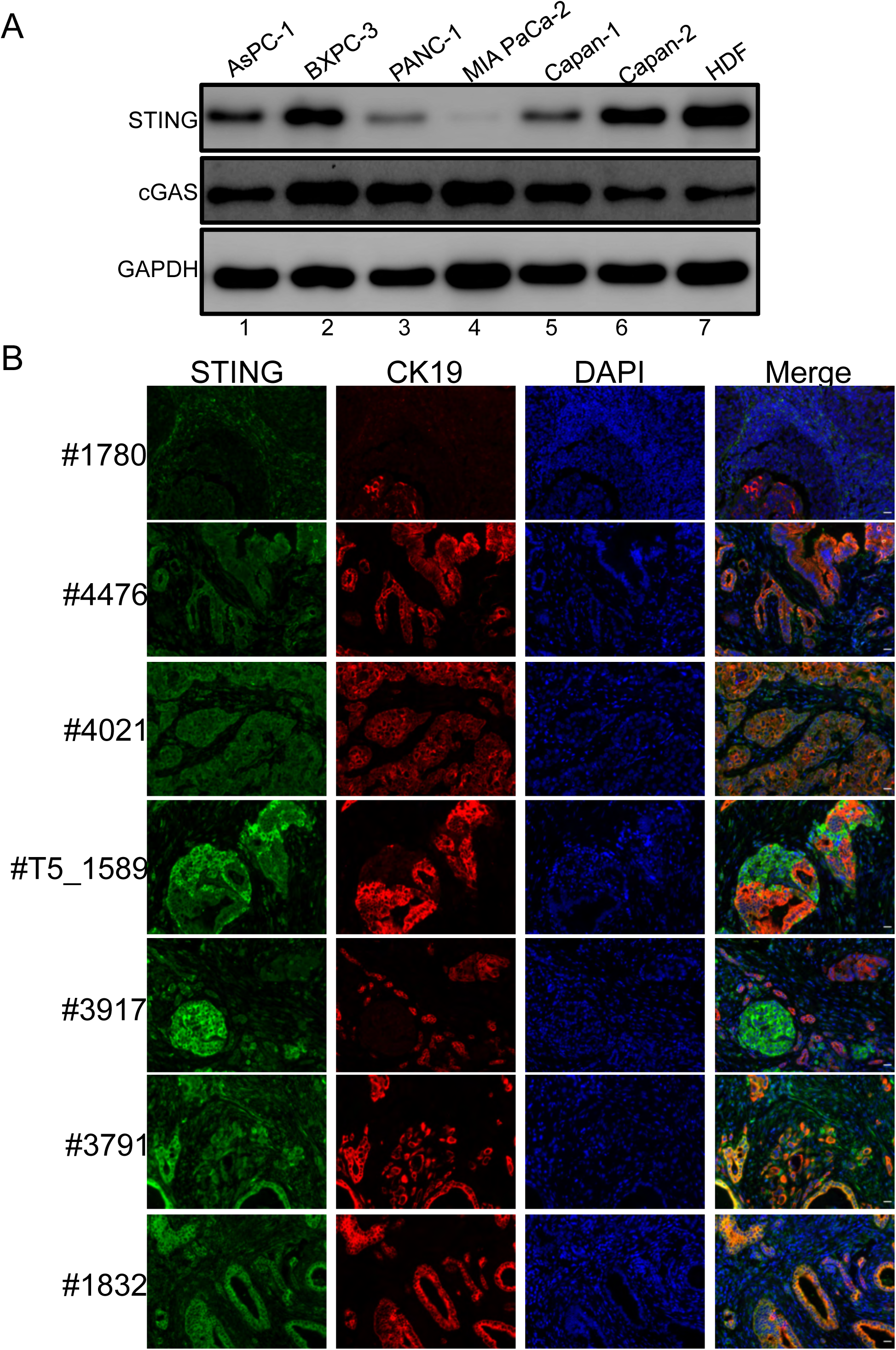
STING is downregulated in PDAC. (A) Whole-cell lysates of PDAC cells and primary HDF cells were immunoblotted using the indicated antibodies. GAPDH was used as a loading control. (B) PDAC lesions were stained for STING (Green) and CK19 (Red), and counterstained with DAPI. Shown are the staining results of pancreatic lesions derived from 7 different patients.

### Identification of a highly active STING gain-of-function mutant

We then set out to establish a new approach for reactivating the STING signaling pathway in STING-silenced cancers using STING gain-of-function genetic mutants. Several single amino acid STING gain-of-function mutants have been identified in autoinflammatory diseases. Among these, the STING^V147L^, STING^N154S^, STING^V155M^, and STING^R284S^ mutants have demonstrated high activity in stimulating downstream innate immune signaling [46,47,60]. Thus, they offer an exciting opportunity to create a simple approach to reactivate the STING signaling pathway. We therefore tested whether these gain-of-function mutants could be used to reignite the anti-tumor activities of the STING signaling pathway in cancer cells. To screen the capability of these STING gain-of-function mutants to inhibit tumor proliferation, we constructed MIA PaCa-2 cells stably expressing either doxycycline (dox)-inducible wild type (WT) STING or one of the STING mutants. Expression of the STING^R284S^ mutant in MIA PaCa-2 cells significantly increased the expression of the early cell death marker cleaved caspase-3 (Fig. 2A) and also inhibited cell proliferation, likely by escalating the number of cells going through cell death (Fig. 2B). In contrast, expression of STING^WT^ and the other STING gain-of-function mutants did not induce such an effect. Notably, all the STING gain-of-function mutants showed lower signal than WT STING (Fig. 2A). This is consistent with previous studies showing that activated STING proteins are quickly degraded [63–65]. Based on the result of this experiment, we selected the STING^R284S^ mutant for our further studies.

**Figure 2.**
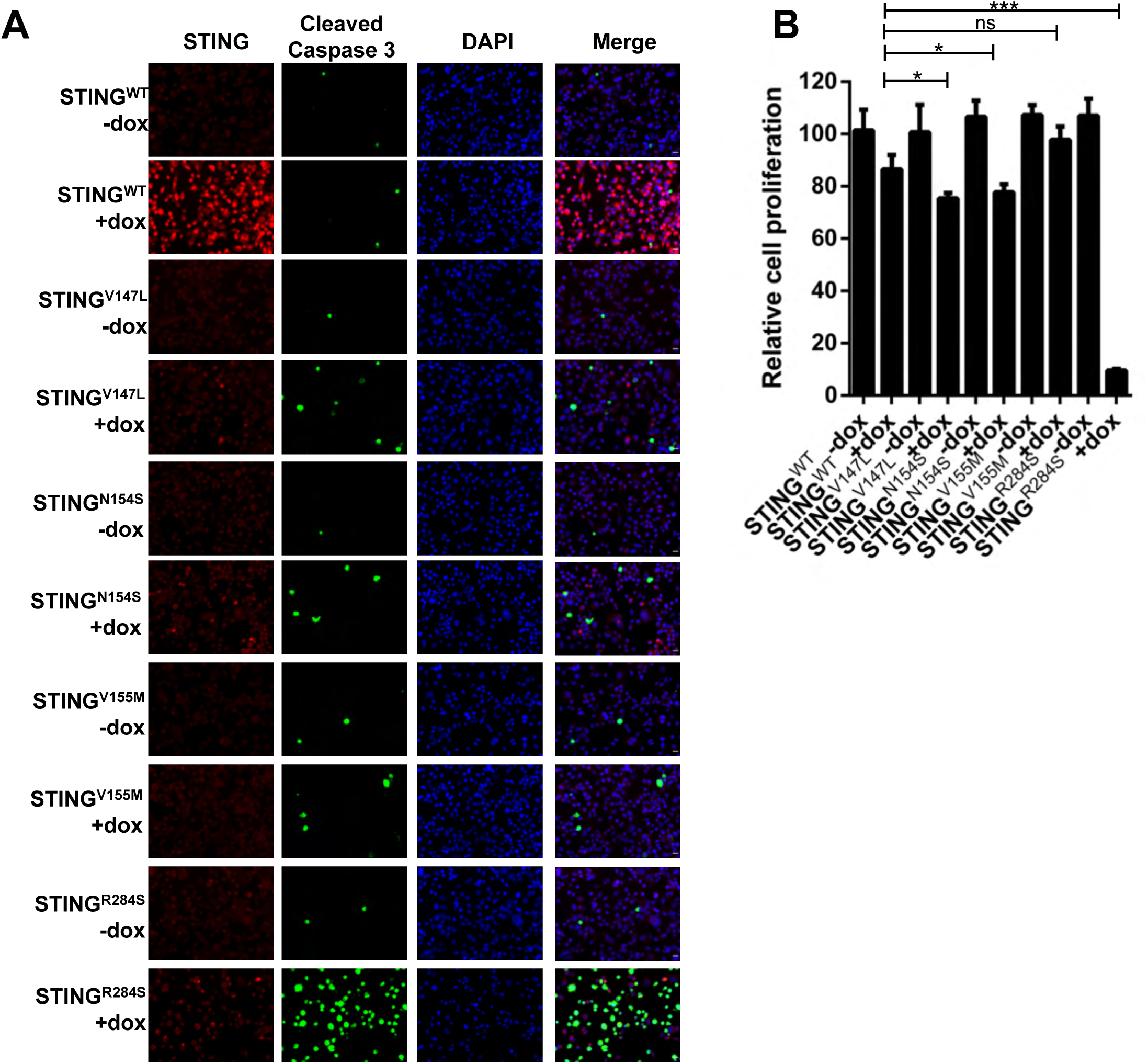
Identification of highly effective STING gain-of-function mutants. (A) MIA PaCa-2 cells stably expressing STING^WT^, STING^V147L^, STING^N154S^, STING^V155M^ or STING^R284S^ were treated with or without 5 µg/mL dox for 48 h. The cells were stained for STING (Red) and Cleaved Caspase-3 (Green). (B) MIA PaCa-2 cells stably expressing STING^WT^, STING^V147L^, STING^N154S^, STING^V155M^, or STING^R284S^ were treated with or without 5 µg/mL dox. At 96 h post-treatment, cell viability was measured by the Titer-GLO 3D cell viability assay (ns: not significant, * P < 0.05, **P < 0.01, ***P < 0.001).

### Ectopic expression of dox-inducible STING^R284S^ induces key anti-tumor cytokine production and cell death in PDAC cells

Our previous study showed that reactivation of the STING signaling pathway not only induces cell death but also generates robust expression of anti-tumor cytokines, such as IFNs, CXCL10, CCL5, and IL6 [14]. To examine whether the STING^R284S^ mutant has the same downstream function, we constructed PDAC cell lines MIA PaCa-2 and BxPC-3 stably expressing dox-inducible STING^WT^ or STING^R284S^. STING expression was efficiently induced by dox treatment in both stable cell lines (Fig. 3A-B, Fig. S1A-B). Compared to STING^WT^, dox-induced STING^R284S^ stimulated the expression of STING downstream anti-tumor cytokines, such as CCL5, CXCL10, IL29, IL6, IFNβ and TNFα (Fig.3C, Fig. S1C). Moreover, compared to un-induced cells and cells expressing dox-induced STING^WT^, expression of STING^R284S^ increased the level of cleaved caspase-3 (Fig. 3A, Fig.S1A) and drastically inhibited the proliferation of these cancer cells (Fig. 3D, Fig. S1D). These results demonstrate that the STING^R284S^ mutant can provoke key anti-tumor cytokine production and cause widespread PDAC cancer cell death. In the *in vivo* setting, tumor cells killed by STING^R284S^ expression could release significant quantities of tumor antigens as well as DNA to activate T cells and amplify both innate and adaptive antitumor responses [66]. Our findings therefore indicate that introducing STING^R284S^ into tumor cells may be a viable therapeutic strategy for treating STING-deficient cancers.

**Figure 3.**
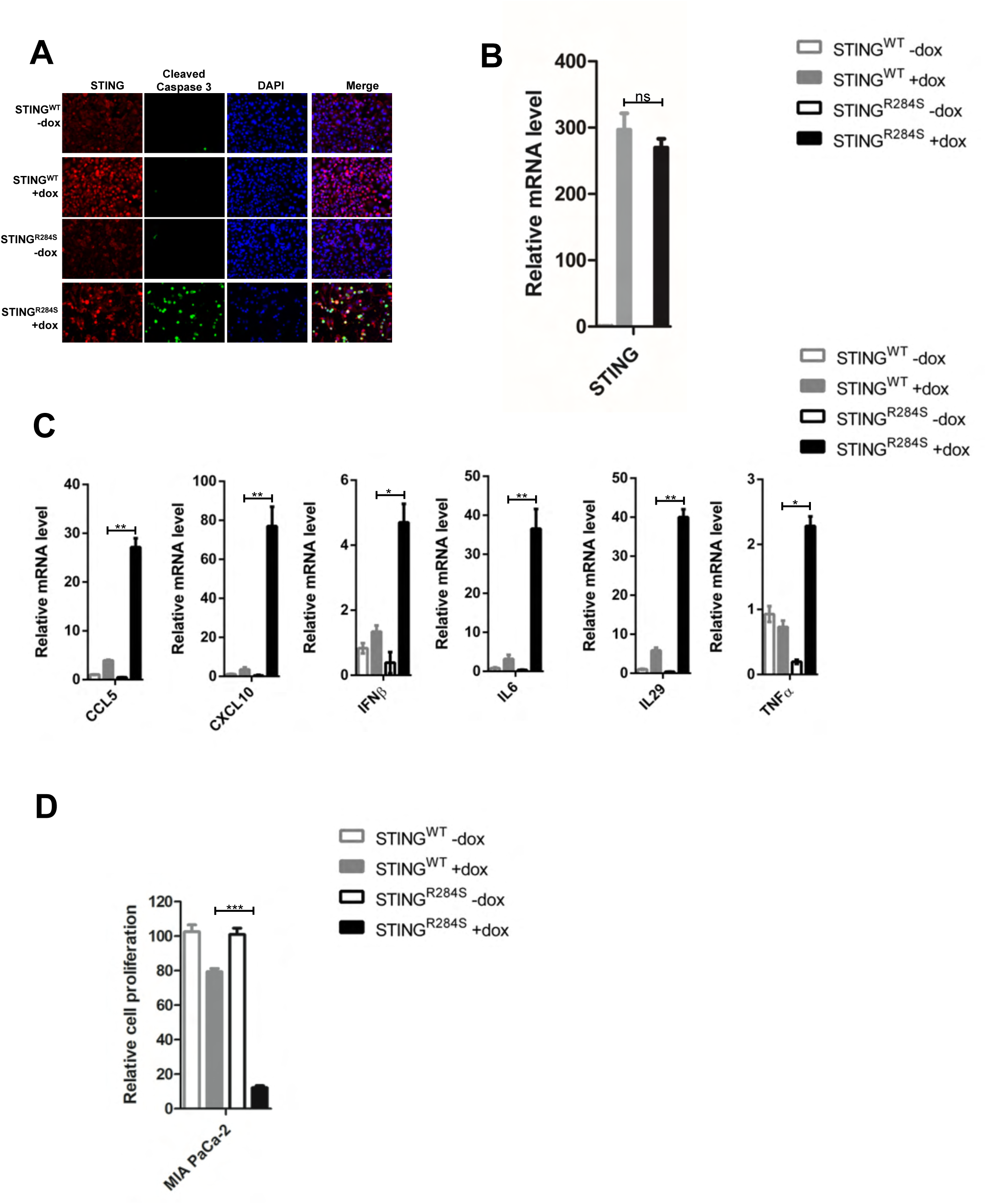
Ectopic expression of dox-inducible STING^R284S^ induces key anti-tumor cytokine production and cell death in PDAC cells. (A-C) MIA PaCa-2 cells stably expressing STING^WT^ or STING^R284S^ were treated with or without 5 µg/mL dox for 48 h. (A) The cells were stained for STING (Red) and Cleaved Caspase-3 (Green). (B) STING^WT^ and STING^R284S^ expression was confirmed by RT-qPCR. (C) The mRNA levels of the indicated genes were measured by RT-qPCR and normalized to GAPDH mRNA levels. The values for untreated STING^WT^ cells were set to 1. (D) MIA PaCa-2 cells stably expressing STING^WT^ or STING^R284S^ were treated with or without 5 µg/mL dox. At 96 h post-treatment, cell viability was measured by the Titer-GLO 3D cell viability assay. Error bars represent SEM of three independent experiments. (ns: not significant, * P < 0.05, **P < 0.01, ***P < 0.001).

### A novel approach to reactivate the STING signaling pathway

We faced a challenge when designing a strategy to deliver the STING^R284S^ mutant into tumor cells as an anticancer therapeutic agent. Viral vectors cannot be used to carry the ‘hot’ STING^R284^ mutant because activation of the STING signaling pathway blocks packaging of many viral-derived vectors [64,67,68]. On the other hand, mRNA-LNP has emerged as a powerful tool for delivering gene expression in cancer cells [69] and also as strong T Helper 1 (Th1) biased adjuvants [70]. Importantly, nucleoside-modified mRNA-LNP can quickly produce abundant protein in target cells while avoiding the host innate immune response [70–81]. Moreover, LNP can be used to package the ‘hot’ STING^R284^ mutant mRNA *in vitro* without activation of the host STING signaling pathway [71, 82]. As a first step to test this strategy, we generated mRNAs encoding STING^R284S^ and STING^WT^ and transfected them into PDAC cells. Compared to mock-transfected cells, robust STING expression was detected in STING^WT^ and STING^R284S^ mRNA-transfected PDAC cells (Fig. 4A-B, Fig. S2A-B). However, only STING^R284S^ mRNA, not STING^WT^ mRNA, stimulated the production of anti-tumor cytokines, such as CCL5, CXCL10, IL29, IL6, IFNβ, and TNFα (Fig. 4C, Fig. S2C). In addition, unlike STING^WT^ mRNA, transfection of STING^R284S^ mRNA significantly elevated the level of cleaved caspase-3 and reduced the cancer cell proliferation rate appreciably (Figs. 4A, 4D, S2A, S2D). These results show that transfection with STING^R284S^ mRNA can specifically stimulate the STING signaling pathway to produce essential anti-tumor cytokines and kill cancer cells.

**Figure 4.**
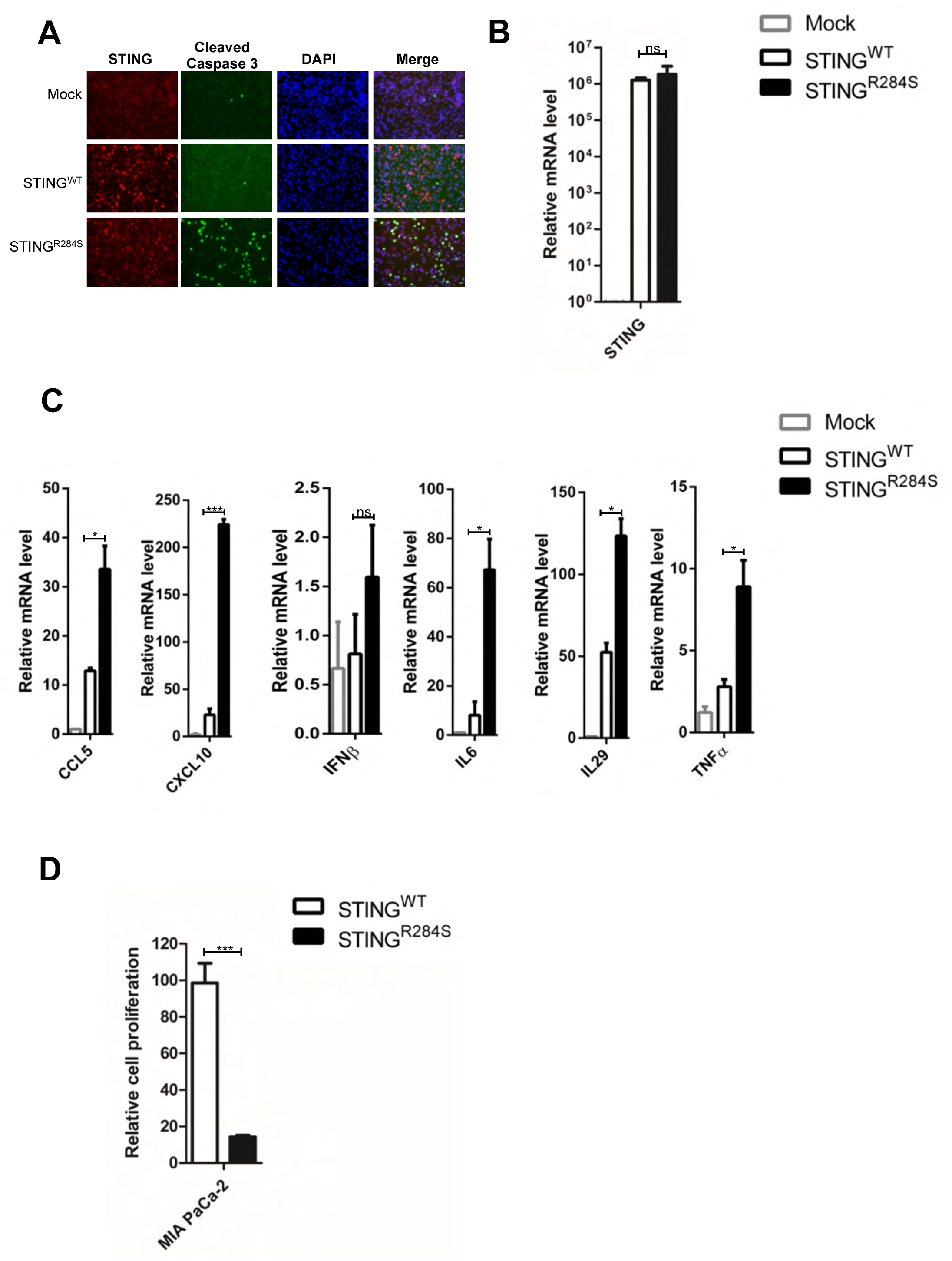
Transfection of STING^R284S^ mRNA activates vital anti-tumor cytokine production and triggers PDAC cell death. (A-C) 10e^4^ MIA PaCa-2 cells were transfected with 0.5 µg STING^WT^ or STING^R284S^ mRNA. At 15 h post-transfection, cells were stained for STING (Red) and Cleaved Caspase-3 (Green) (A), STING^WT^ and STING^R284S^ expression was confirmed by RT-qPCR (B), and the mRNA levels of the indicated genes were measured by RT-qPCR and normalized to the GAPDH mRNA level (C). The values for untreated cells (Mock) were set to 1. (D) 0.5×10e^4^ MIA PaCa-2 cells were transfected with 1 µg STING^WT^ or STING^R284S^ mRNA. At 15 h post-transfection, cell viability was measured by the Titer-GLO 3D cell viability assay. Error bars represent SEM of three independent experiments. (ns: not significant, * P < 0.05, **P < 0.01, ***P < 0.001).

### STING^R284S^ expression delivered by mRNA-LNP activates vital anti-tumor cytokines and induces PDAC cell death

To further develop a therapeutic approach, we tested whether LNP can be used to deliver the STING^R284S^ mRNA into cancer cells. The LNP we exploited in this study have been shown to efficiently deliver genes *in vivo* [83]. However, we did not observe significant expression of STING^WT^ and STING^R284S^ in MIA PaCa-2 and BxPC-3 cells treated with the respective mRNA-LNP (Figs. 5A and S3A, rows 2, 5). We reasoned that this could be due to a lack of Human Apolipoprotein E (APOE) in our *in vitro* cultures. In the *in vivo* setting, APOE plays an important role in the cellular uptake of physiological lipoproteins through binding to low-density lipoprotein (LDL) receptors [84, 85]. When mixed with mRNA-LNP before transduction, human APOE4 has been shown to radically increase mRNA-LNP transduction efficiency *in vitro* [84, 85]. We therefore tested whether mixing STING^WT^ or STING^R284S^ mRNA-LNP with APOE4 could facilitate the delivery of mRNA into PDAC cells. We found that APOE4 robustly stimulates the delivery of mRNA-LNP into PDAC cells in a dose-dependent manner (Fig. 5A, Fig. S3A). Compared to untreated cells, higher levels of STING^WT^ and STING^R284S^ mRNA were detected by RT-PCR in cells treated with a combination of mRNA-LNP and APOE4 (Fig. 5B, Fig. S3B). The combined treatment of STING^R284S^ mRNA-LNP and APOE4 also significantly augmented the expression of key anti-tumor cytokines, such as CCL5, CXCL10, IL29, IL6, and TNFα, as compared with treatment using only STING^WT^ mRNA (Fig.5C, Fig. S3C). Moreover, LNP-delivered STING^R284S^ mRNA not only induced the production of cleaved caspase-3 in PDAC cells, but also significantly inhibited the proliferation of these cells (Figs. 5A, 5D, S3A, S3D). Importantly, the same treatment did not repress the proliferation of CD8+ T cells (Fig. S3E) (see discussion). These results demonstrate that human APOE4 can efficiently promote the delivery of mRNA-LNP into target cells, allowing the robust expression of ‘hot’ STING^R284S^ to induce essential anti-tumor cytokines and eradicate cancer cells.

**Figure 5.**
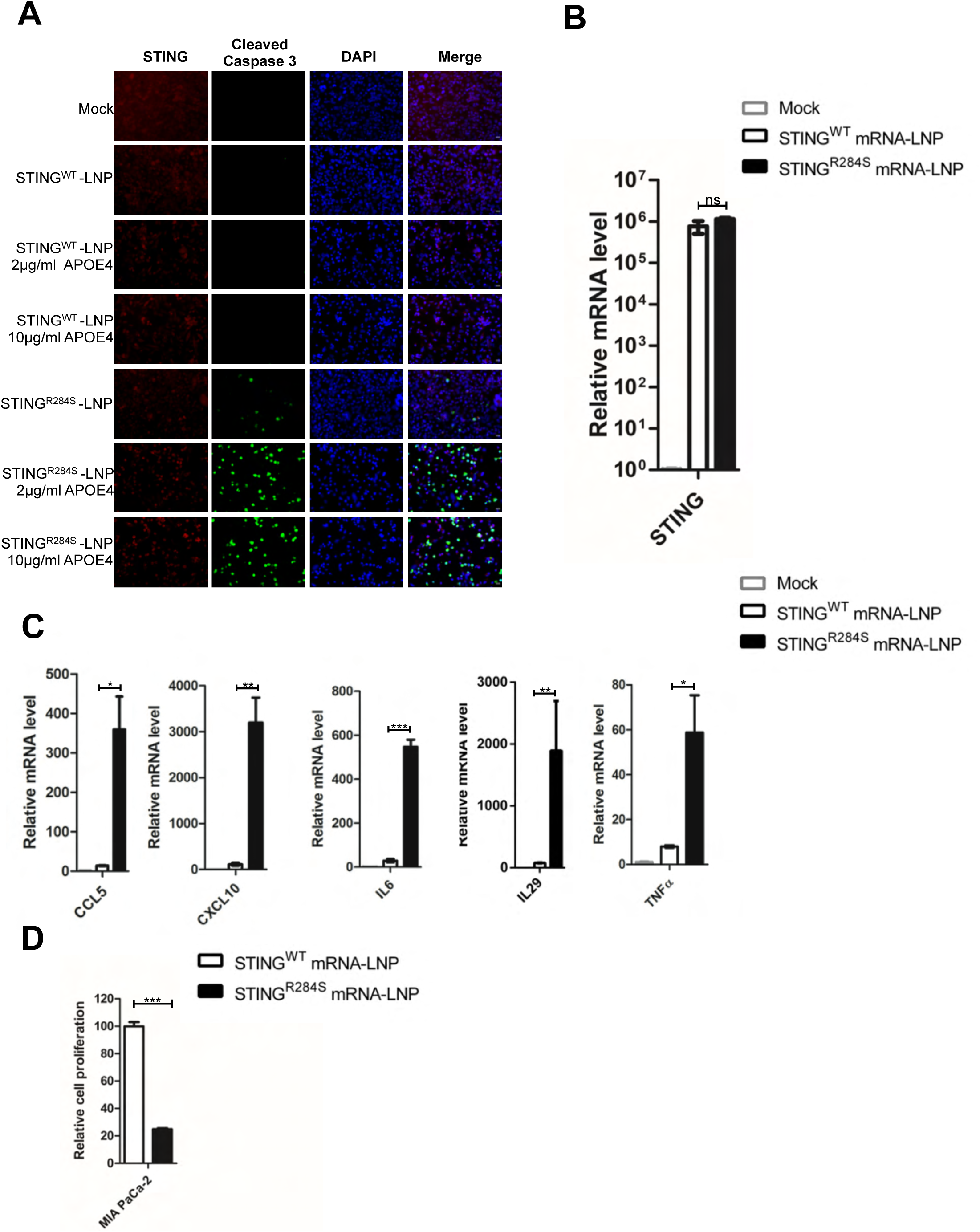
STING^R284S^ delivered by mRNA-LNP activates essential anti-tumor cytokine production and kills PDAC cells. (A) 2×10e^4^ MIA PaCa-2 cells were treated with 1 µg STING^WT^ or STING^R284S^ mRNA-LNP, which were pre-mixed with the indicated concentration of the recombinant human APOE4 protein. At 16 h post-transfection, cells were stained for STING (Red) and Cleaved Caspase-3 (Green). (B-C) 10e^4^ MIA PaCa-2 cells were treated as in (A) using 10 µg/ml human APOE4 protein. At 16 h post-transfection, STING^WT^ and STING^R284S^ expression was confirmed by RT-qPCR (B), and the mRNA levels of the indicated genes were measured by RT-qPCR and normalized to the GAPDH mRNA level (C). The values for untreated cells (Mock) were set to 1. (D) 0.5×10e^4^ MIA PaCa-2 cells were treated as in (B). At 16 h post-transfection, cell viability was measured by the Titer-GLO 3D cell viability assay. Error bars represent SEM of three independent experiments. (ns: not significant, * P < 0.05, **P < 0.01, ***P < 0.001).

### STING^R284S^ mRNA-LNP also trigger vital anti-tumor cytokine production and cell death in MCC cells

We recently found that STING is also silenced in some MCC tumors [14]. 80% of MCCs have integrated Merkel cell polyomavirus (MCPyV) genomes [86]. Our previous studies showed that STING is specifically repressed in MCPyV^+^ MCC cell lines [14]. By analyzing published RNA-seq data [87], we discovered that while STING is amply expressed in the MCPyV^-^ MCC cell line UISO, STING RNA expression levels are nearly undetectable in all six classic MCPyV^+^ MCC cell lines: MKL-1, MKL-2, MS-1, WaGa, PeTa, and BroLi (Fig.S4A). The RNA-seq data also indicated that when compared with other MCPyV^+^ MCC cell lines, STING RNA expression is slightly higher in PeTa cells (Fig.S4A) [87]. However, Western-blot analysis reveals that, similar to MKL-1 cells, STING protein expression in PeTa cells is completely imperceptible (Fig. S4B). This study therefore confirmed that STING expression is suppressed in all of the classic MCPyV^+^ MCC cell lines we have examined.

Encouraged by the observed antitumor activity of the STING^R284S^ mRNA-LNP in PDAC cells, we tested whether our LNP approach could be applied to stimulate the same positive response in MCC cells. We first optimized the mRNA-LNP delivery conditions for MCC cells using firefly luciferase mRNA-LNP. We ascertained that 10 ug/ml of human APOE4 was also the ideal concentration for delivering mRNA-LNP into MCC cells (Fig. S5). When compared with the untreated MKL-1 and MS-1 MCC cells, robust STING expression was detected in both STING^WT^ and STING^R284S^ mRNA-LNP-treated cells (Figs. 6A-B, S6A-B). However, only delivery of STING^R284S^ mRNA-LNP, and not STING^WT^ mRNA-LNP, stimulated expression of the key anti-tumor cytokines CCL5, CXCL10, IL29, IL6, IFNβ, and TNFα (Figs. 6C, S6C). Compared to STING^WT^ mRNA-LNP, treating MCC cells with STING^R284S^ mRNA-LNP also elevated the level of cleaved caspase-3 and greatly inhibited cell proliferation (Figs. 6A, 6D, S6A, S6D). These results demonstrated that the STING^R284S^ mRNA-LNP could also induce anti-tumor cytokine expression and cell death in tested MCC cell lines.

**Figure 6.**
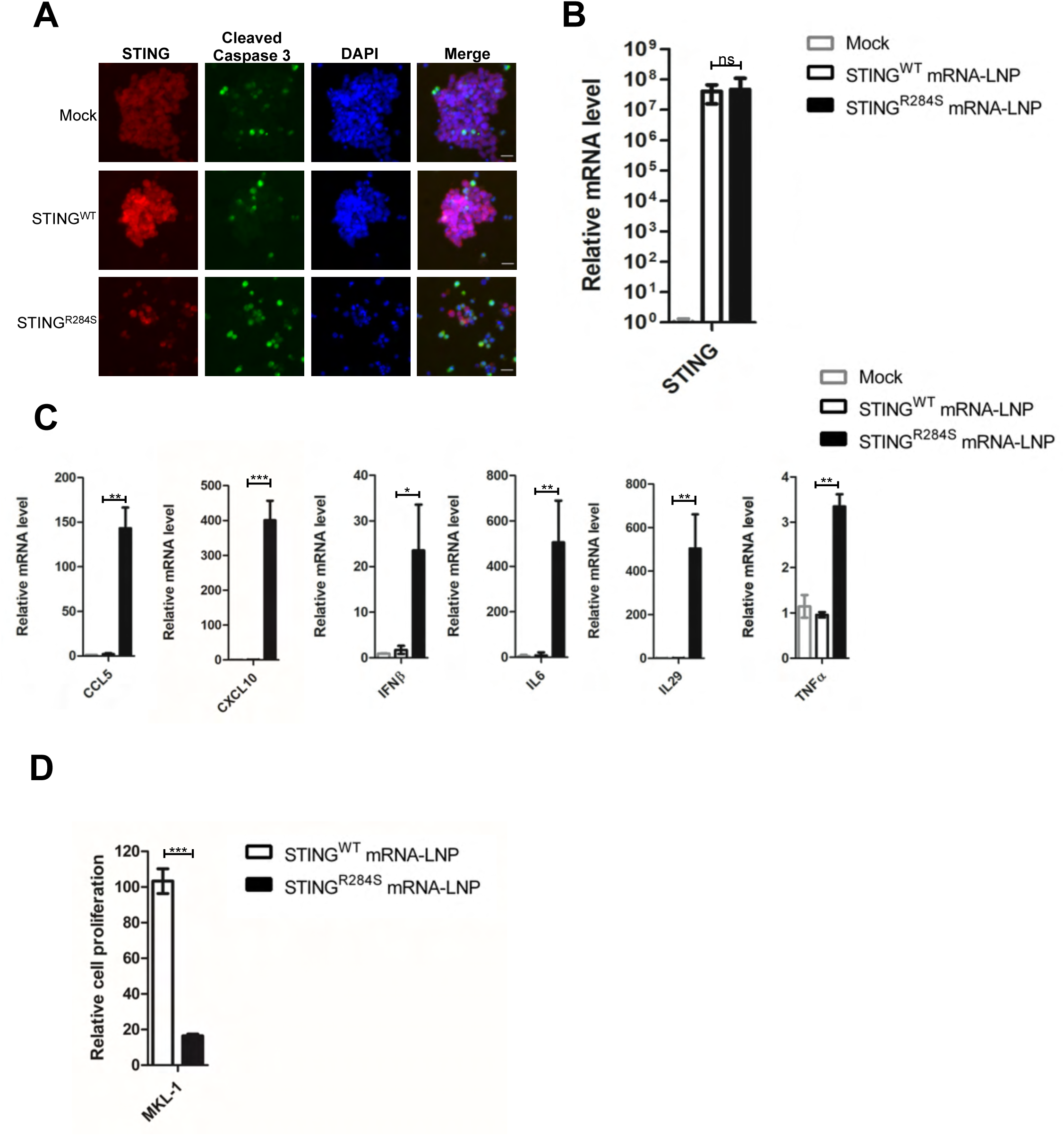
STING^R284S^ mRNA-LNP can trigger vital anti-tumor cytokine production and cell death in MCC cells. (A) 10e^4^ MKL-1 cells were treated with 1 µg STING^WT^ or STING^R284S^ mRNA-LNP, which were pre-mixed with the indicated concentration of the recombinant human APOE4 protein. At 16h post-transfection, cells were stained for STING (Red) and Cleaved Caspase-3 (Green). (B-C) 10e^4^ MKL-1 cells were treated as in (A) using 10 µg/ml human APOE4 protein. At 16h post-transfection, STING^WT^ and STING^R284S^ expression was confirmed by RT-qPCR (B), and the mRNA levels of the indicated genes were measured by RT-qPCR and normalized to the GAPDH mRNA level (C). The values for untreated cells (Mock) were set to 1. (D) 0.5×10e^4^ MKL-1 cells were treated as in (B) at 0 and 24 h. At 40 h post-transfection, cell viability was measured by the Titer-GLO 3D cell viability assay. Error bars represent SEM of three independent experiments. (ns: not significant, * P < 0.05, **P < 0.01, ***P < 0.001).

In summary, we demonstrated that STING^R284S^ mRNA-LNP robustly activate the STING signaling pathway in cancer cells, leading to the production of key anti-tumor cytokines as well as cancer cell death. Therefore, STING^R284S^ mRNA-LNP could be exploited as a promising anticancer drug for treating STING-deficient cancers.

## Discussion

Currently, several therapeutic approaches such as PD-1/PD-L1 and CTLA-4 inhibitors have been appraised in attempts to combat aggressive cancers such as PDACs and MCCs, but have failed to produce durable responses in PDACs [88] and led to treatment resistance in some MCCs [89]. Therefore, alternative therapeutics are still needed for treating these lethal cancers.

The complex tumor microenvironment presents a major barrier to developing broadly effective therapies. The TME of PDAC is known to be immunosuppressive. Although tumor infiltration of T lymphocytes was positively correlated with overall patient survival [90], the PDAC TME has very few tumor-infiltrating CD8+ cytotoxic T cells and CD4+ T helper cells, and instead exhibits an increased presence of regulatory T cells, tumor-associated macrophages, and myeloid-derived suppressor cells [91, 92]. We and others have reported that the STING signaling pathway is dysfunctional in several cancers [14,20,43]. Thus, we examined the expression of key components of this pathway, cGAS and STING, in PDAC cell lines. We found that all tested pancreatic cancer cell lines maintain highly expressed cGAS, but STING is significantly downregulated in many of the PDAC cell lines and tissues (Fig. 1). In light of the STING function in stimulating antitumor response, we speculated that STING repression might contribute to the immunosuppressive TME of PDACs and that reactivating STING might represent a viable strategy for heating up the immunologically ‘cold’ TME in PDAC.

To stimulate STING activity in PDAC cells, we first screened several ‘hot’ STING mutants. We discovered that only the STING^R284S^ mutant, but not STING^WT^ nor the other STING gain-of-function mutants such as STING^V147L^, STING^N154S^, and STING^V155M^, could specifically inhibit the growth of STING-silenced MIA PaCa-2 cells (Fig. 2). The result correlates appropriately with the clinical impact of these gain-of-function mutations. For example, the STING^V147L^, STING^N154S^ and STING^V155M^ mutants were identified in patients who died at an age of at least 9 years [49], but the STING^R284S^ mutant was derived from a patient who died at approximately 9 months of age [47]. Together, our finding suggests that, among all of the mutants tested, STING^R284S^ has the highest activity in stimulating the STING signaling pathway. This discovery provides the molecular basis for using the STING^R284S^ mutant to develop STING-targeted immunotherapies.

Our further studies demonstrated that STING^R284S^ mRNA-LNP could be efficiently delivered into PDAC cells to induce cytokines/chemokines crucial for promoting intratumoral infiltration of CD8^+^ T cells. More importantly, STING^R284S^ expression also induces robust cell death in STING-silenced cancers (Figs. 5 and S3). MCCs also have an immunologically ‘cold’ TME and STING is invariably repressed in the MCPyV^+^ MCC tumors we have examined. We further demonstrated that STING^R284S^ mRNA-LNP could also be utilized to activate STING downstream antitumor activity in MCC tumor cells. In summary, by harnessing the hyperactive immune-stimulatory activity of the STING^R284S^ mutant and the delivery capability of mRNA-LNP, we have provided evidence for using the naturally occurring STING^R284S^ mutant as a novel therapeutic tool to reactivate the antitumor response in the immunologically ‘cold’ pancreatic cancer and in other STING-silenced tumors.

Several observations suggest that STING^R284S^ mRNA-LNP hold great promise for developing a cancer immunotherapy. First, when compared with wild type STING, ‘hot’ STING mutants such as STING^R284S^ are more responsive to cGAMP [46,47,49,51]. When delivered into tumor cells by mRNA-LNP, STING^R284S^ can be further activated by the large amount of damaged DNA present in these cells, spurring robust antitumoral activity. Therefore, no additional STING agonist is needed to stimulate ‘hot’ STING mutants, increasing the feasibility for clinical application. Secondly, pancreatic cancers possess few tumor-specific new epitopes (neoantigens) [12]. STING^R284S^ mRNA-LNP-induced cell death will play a crucial role in exposing neoantigens of tumors to the host immune system. The large amount of tumor antigens released by the dead cells can be engulfed by antigen-presenting cells (APCs) and presented to T cells to generate systemic antitumor immunity and amplify the tumoricidal effect. This process could also induce adaptive antitumor immunity for rejecting distant metastases and providing long-living immunologic memory. Thirdly, STING^R284S^-mediated cell death can also directly reduce cancer burden, which is also clearly beneficial to cancer immunotherapy [93, 94]. Finally, mRNA-LNP has an intrinsic adjuvant effect that can stimulate T follicular helper cells (Tfh) responses and promote the production of effective CD8^+^ T cells [70,78,95]. An additional advantage is that multiple mRNAs can be combined together or with other drugs to be encapsidated into LNP [78-81,96].

The STING^R284S^ mRNA-LNP approach can be used to restore STING expression and function in STING-deficient tumors in order to stimulate anti-tumor immune responses and directly kill the tumor cells (Figs 4C, 5C, S3C, and S5C). Anti-tumor cytokines have safety concerns when systemically administered; however, gene expression driven by intratumorally-injected mRNA-LNP has been detected mainly in the tumor sites but not in major vital organs [96, 97]. Therefore, local delivery using STING^R284S^ mRNA formulated in LNP could overcome the specificity issue and reveal a safe approach to leverage the cytokine effects [98, 99]. Additionally, overstimulation of STING in T cells could introduce cell death and cytotoxicity, which counteracts the desired antitumor immune response [17,55,100–103]. Interestingly, we found that while STING^R284S^ mRNA-LNP can effectively repress cancer cell proliferation, it does not inhibit the growth of CD8^+^ T cells (Figs. 4D, 5D, S3D, S5D and S3E). This is consistent with previous studies confirming that T cells are not susceptible to transfection by exogenous mRNA delivered in LNP [104]. Therefore, mRNA-LNP-mediated intratumoral delivery of STING^R284S^ will allow specific activation of STING signaling in tumor tissues without introducing antiproliferative effects in lymphocytic immune cells. Because mRNA-LNP delivery is transient, it also allows for greater control of the treatment process. So far, all STING^R284S^ mRNA-LNP studies were performed *in vitro.* Plans are underway to establish STING-negative tumor models in mice, which will be used to examine the efficacy of the STING^R284S^ mRNA-LNP in stimulating T cell intratumoral infiltration and killing of tumor cells *in vivo*. Furthermore, we are also developing specific targeting strategies in order to apply the STING^R284S^ mRNA-LNP for treating metastatic disease.

STING agonists are being actively pursued as new cancer immunotherapies [36,44,45,105], but few have generated positive clinical outcome [39, 40]. As shown by our group and others, STING is silenced in many cancers [14,20,43]. Our findings could explain why traditional STING agonists will not work in STING-silenced cancers, as the antitumor efficacy of these agonists obligatorily depends on STING expression to begin with [44]. When delivered into noncancerous cells, the classic STING agonists can also induce inflammatory diseases and cancers [17, 103]. Our STING^R284S^ mRNA-LNP approach therefore represents a novel therapeutic strategy that can overcome the limitations and toxicity of conventional STING agonist-based therapies. It also possesses broader potential for overcoming the immunosuppressive microenvironment in other STING-deficient ‘cold’ tumors.

## Materials and methods

### Cell culture and cancer lesions

Primary foreskin dermal fibroblasts [106], human embryonic kidney 293T (HEK293T), MIA PaCa-2, and PANC-1 cells were grown in Dulbecco’s modified Eagle’s medium supplemented with 10% fetal calf serum. BxPC-3 and AsPC-1 cells were grown in RPMI 1640 medium supplemented with 10% fetal calf serum. Capan-1 and Capan-2 cells were grown in McCoy’s 5A Medium supplemented with 10% fetal calf serum. MKL-1 and MS-1 cells were grown in RPMI 1640 medium supplemented with 20% fetal calf serum. All the cells were incubated at 37°C in humidified air containing 5% CO2. Primary CD8^+^ T cells from healthy donors were provided by the Human Immunology Core at the University of Pennsylvania. These cells were grown in RPMI 1640 medium supplemented with 10% heat-inactivated FBS, L-glutamine, IL-2 and Penicillin-Streptomycin. PDAC tissues were obtained from the Tumor Tissue and Biospecimen Bank at the University of Pennsylvania.

### Western blot analysis

To prepare whole cell lysates, cells were lysed in lysis buffer (10 mM HEPES, pH 7.9, 500 mM NaCl, 3 mM MgCl_2_, 1 mM DTT, 1 mM PMSF, 0.5% Triton X-100 supplemented with protease inhibitors). After 30 minutes of incubation on ice, whole cell lysates were centrifuged at 15,000g for 10 min at 4°C to remove the debris. Protein concentrations were determined using the Bradford assay. The protein samples were resolved on SDS-PAGE gels, transferred onto PVDF membranes, and immunoblotted with specific primary antibodies as indicated in the figure legends. The primary antibodies used in this study include anti-STING (1:2000, 13647S, Cell Signaling Technology), anti-cGAS (1:1000, 15102, Cell Signaling Technology), and anti-GAPDH (1:2000, 5174S, Cell Signaling Technology). The secondary antibody used was HRP-linked anti-rabbit IgG (1:3000, 7074S, Cell Signaling Technology). Western blots were developed using SuperSignal™ West Pico PLUS Chemiluminescent Substrate (Thermo Scientific) and images were captured using a GE imaging system.

### Cell proliferation assay

Cell viability was measured with CellTiter-Glo 3D (Promega) following the manufacturer’s instructions [107].

### Reverse transcription and quantitative real-time PCR

Total RNA was isolated using NucleoSpin RNA II Kit (Macherey-Nagel) in pursuance of the manufacturer’s protocol. Reverse transcription (RT) was performed using a 20 µl reaction mixture containing 350 ng of total RNA, random hexamer primers (Invitrogen), dNTPs (Invitrogen), and M-MLV reverse transcriptase (Invitrogen). Quantitative real-time PCR (qPCR) was performed using a CFX96 real-time PCR detection system (Bio-Rad) with IQ SYBR Green supermix (Bio-Rad). Primer sequences are the same as listed in [14]. The mRNA level of each gene was normalized to the GAPDH mRNA level.

### Immunofluorescent staining

Cells were fixed with 3% paraformaldehyde in PBS for 20 minutes. Immunofluorescent (IF) staining was performed as previously described [108]. The following primary antibodies were used: anti-CK19 (1:200, 4558, Cell Signaling Technology), anti-STING (1:500 for cell staining, 1:20 for tissue staining, 19851-1-AP, Proteintech), and anti-Cleaved Caspase-3 (Asp175) (1:500, 9661, Cell Signaling Technology). The secondary antibodies used were Alexa Fluor 594 goat anti-mouse IgG (1:500, A-11032, ThermoFisher Scientific) and Alexa Fluor 488 goat anti-rabbit IgG (1:500, A-11008, ThermoFisher Scientific). All IF images were collected using an inverted fluorescence microscope (IX81; Olympus) connected to a high-resolution charge-coupled-device camera (FAST1394; QImaging). Images were analyzed and presented using SlideBook (version 5.0) software (Intelligent Imaging Innovations, Inc.). The scale bars were added using ImageJ software.

### Recombinant plasmid construction

pTEV-STING^WT^-A101 plasmid containing codon-optimized STING^WT^ gene was synthesized by Genewiz. pTEV-STING^R284S^-A101 plasmid containing the STING^R284S^ gene was generated from codon-optimized human STING^WT^ by PCR-based site-directed mutagenesis.

### mRNA production

Using linearized plasmid pTEV-STING^WT^-A101 and pTEV-STING^R284S^-A101, the STING mRNA was produced with T7 RNA polymerase. During mRNA synthesis, 1-methylpseudouridine-5′-triphosphate (TriLink) was used instead of UTP to generate modified nucleoside-containing mRNA. The STING mRNA was co-transcriptionally capped using CleanCap (TriLink) and purified as described previously [70]. The STING mRNA was analyzed by agarose gel electrophoresis and stored frozen at −80 °C.

### mRNA transfection

Transfection of human pancreatic MIA PaCa-2 and BxPC-3 cells was performed with TransIT-mRNA (Mirus Bio) according to the manufacturer’s instructions. Specifically, mRNA (1 µg) was combined with TransIT-mRNA reagent (3 µl) and boost reagent (3 µl) in 100 µl of serum-free medium, and the complex was added to 10e^5^ cells in 500 µl complete medium. Cells were harvested at 15-16 h after transfection.

### LNP encapsulation of the mRNA

Purified STING mRNAs were encapsulated in LNP using a self-assembly process in which an aqueous solution of mRNA at pH 4.0 is rapidly mixed with a solution of lipids dissolved in ethanol. LNP used in this study were similar in composition to those described previously [71], which contain an ionizable cationic lipid (proprietary to Acuitas), phosphatidylcholine, cholesterol and PEG-lipid. The ionizable cationic lipid and LNP composition are described in the patent application WO 2017/004143. The diameter and polydispersity index of LNP was measured by dynamic light scattering using a Zetasizer Nano ZS instrument (Malvern Instruments Ltd) and an encapsulation efficiency of ∼95% as determined using a Ribogreen assay. RNA-LNP formulations were stored at - 80 °C at an RNA concentration of ∼1 µg/µl.

### Mutagenesis Primers

The sequences for the primers used in STING mutagenesis are:

STING^V155M^F: AACATGGCCCATGGGCTGGCATGG
STING^N154S^F: AGCGTGGCCCATGGGCTGGCATGG
STING^N154S^R: GAAATTCCCTTTTTCACACACTGCAGAG
STING^V147L^F: CTGTGTGAAAAAGGGAATTTCAACGTGG
STING^V147L^R: TGCAGAGATCTCAGCTGGGG

### Statistical analyses

Statistical analysis was performed using the unpaired t-test of GraphPad Prism software (Version 7.0) to compare the data from the control and experimental groups. A two-tailed P value of <0.05 was considered statistically significant.

## Acknowledgments

We thank the Human Immunology Core (University of Pennsylvania) through Grants P30-CA016520 and P30AI045008 for providing purified human CD8^+^ T cells; Dr. Erle S. Robertson (University of Pennsylvania) for pancreatic cancer cell lines MIA PaCa-2, BxPC-3, PANC-1, AsPC-1, Capan-1, and Capan-2; the Tumor Tissue and Biospecimen Bank (University of Pennsylvania) for providing PDAC tissues; Dr. Dominic De Nardo (Monash University) for pTRIPZ-STING^WT^ and pTRIPZ-STING^R284S^ plasmids; and Drs. Ben Stanger and Ranran Wang for helpful discussion.

## Author Contributions

Wei Liu: conceptualization, formal analysis, investigation, methodology, visualization, writing - original draft, writing- review & editing; Mohamad-Gabriel Alameh: resources, writing- review & editing; June F. Yang: methodology, writing - original draft, writing- review & editing; Jonathan R. Xu: literature research, conceptualization, writing- review & editing; Paulo JC Lin: resources, writing- review & editing; Ying K Tam: resources, writing- review & editing; Drew Weissman: supervision, resources, writing- review & editing; Jianxin You: conceptualization, funding acquisition, project administration, supervision, writing- original draft, writing- review & editing.

---

**Figure S1.**
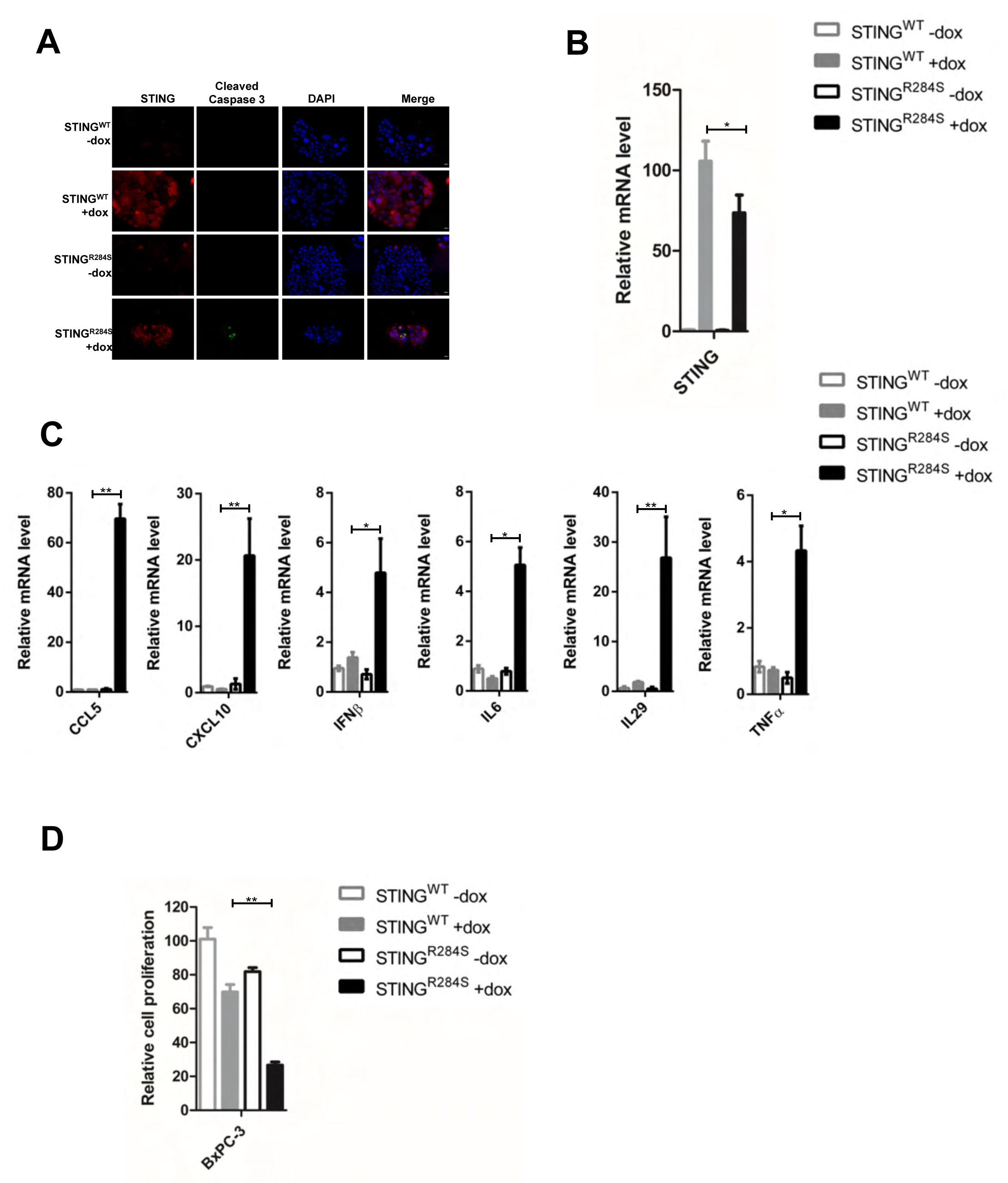
Ectopic expression of dox-inducible STING^R284S^ stimulates key anti-tumor cytokine production and cancer cell death in PDAC cells. (A-C) BxPC-3 cells stably expressing STING^WT^ or STING^R284S^ were treated with or without 5µg/mL dox for 24 h. (A) The cells were stained for STING (Red) and Cleaved Caspase-3 (Green). (B) STING^WT^ and STING^R284S^ expression was confirmed by RT-qPCR. (C) The mRNA levels of the indicated genes were measured by RT-qPCR and normalized to the GAPDH mRNA level. The values for untreated STING^WT^ cells were set to 1. (D) BxPC-3 cells stably expressing STING^WT^ or STING^R284S^ were treated with or without 5µg/mL Dox. At 72 h post-treatment, cell viability was measured by the Titer-GLO 3D cell viability assay. Error bars represent SEM of three independent experiments. (ns: not significant, * P < 0.05, **P < 0.01, ***P < 0.001).

**Figure S2.**
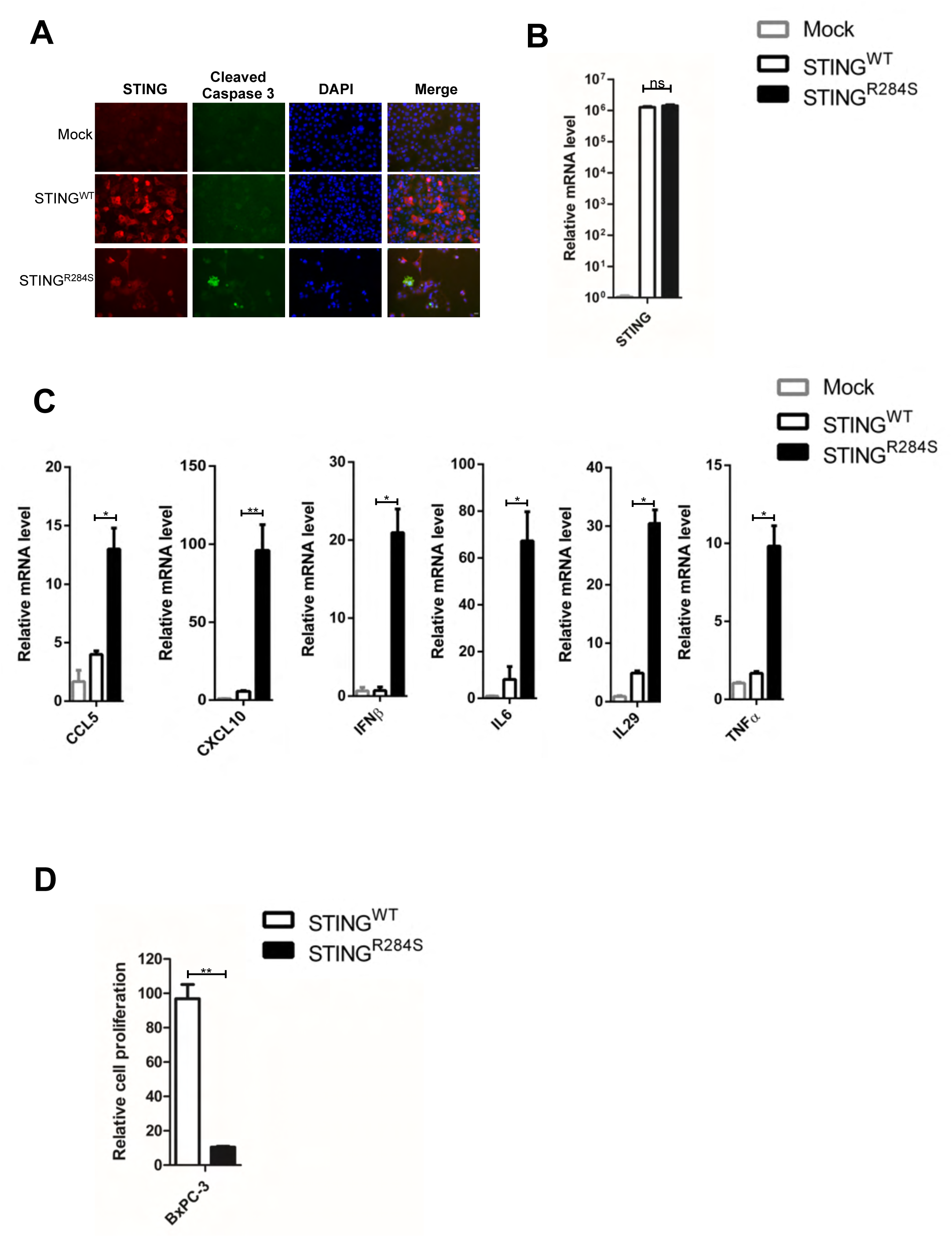
Transfection of STING^R284S^ mRNA activates vital anti-tumor cytokine production and triggers PDAC cell death. (A-C) BxPC-3 cells were transfected with 0.5 µg STING^WT^ or STING^R284S^ mRNA. At 15 h post-transfection, cells were stained for STING (Red) and Cleaved Caspase-3 (Green) (A), STING^WT^ and STING^R284S^ expression was confirmed by RT-qPCR (B), and the mRNA levels of the indicated genes were measured by RT-qPCR and normalized to the GAPDH mRNA level (C). The values for untreated cells (Mock) were set to 1. (D) BxPC-3 cells were transfected with 1 µg STING^WT^ or STING^R284S^ mRNA. At 15 h post-transfection, cell viability was measured by the Titer-GLO 3D cell viability assay. Error bars represent SEM of three independent experiments. (ns: not significant, * P < 0.05, **P < 0.01, ***P < 0.001).

**Figure S3.**
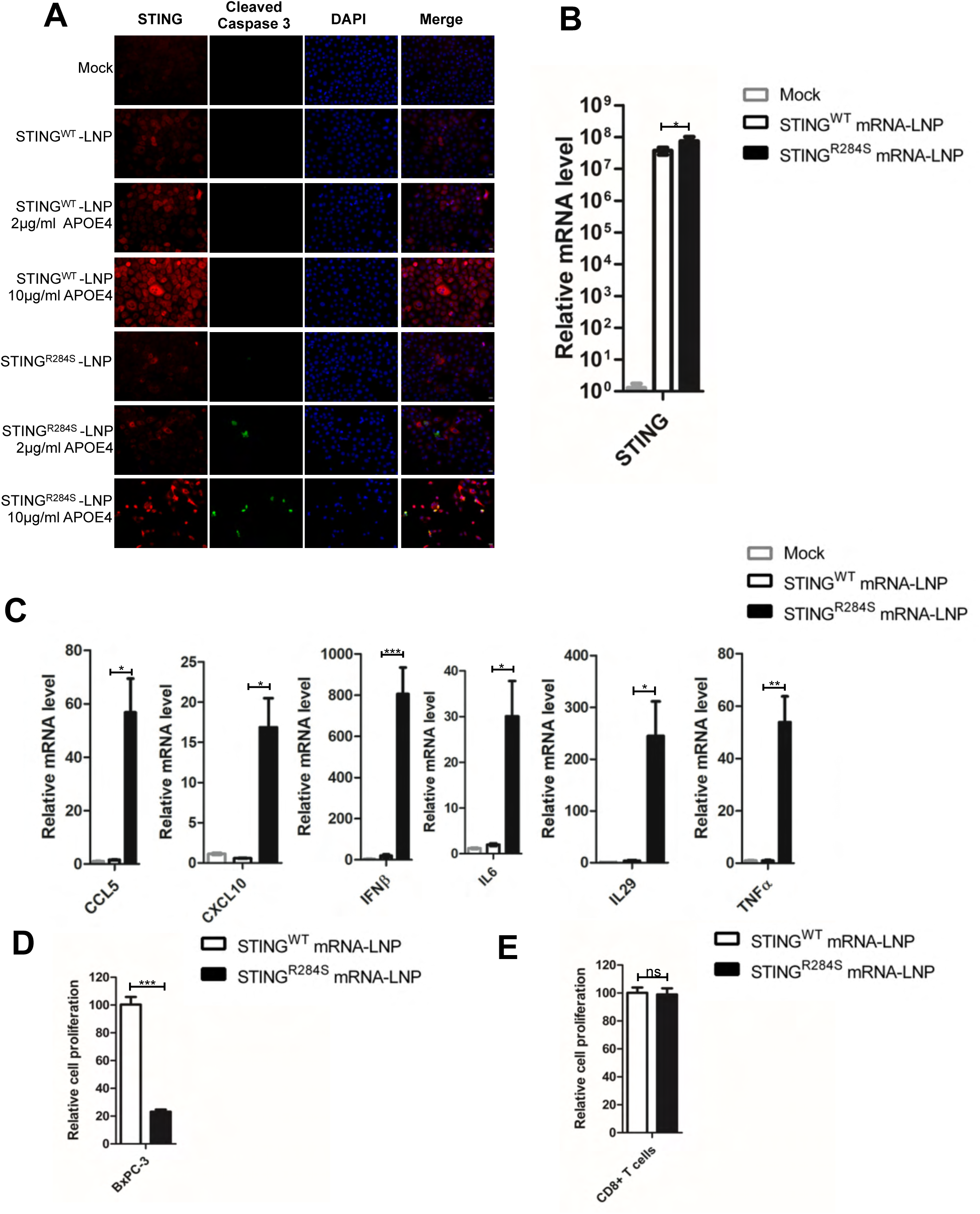
STING^R284S^ delivered by mRNA-LNP activates essential anti-tumor cytokine production and kills PDAC cells. (A) 10e^4^ BxPC-3 cells were treated with 1 µg STING^WT^ or STING^R284S^ mRNA-LNP, which were pre-mixed with the indicated concentration of the recombinant human APOE4 protein. At 16h post-transfection, cells were stained for STING (Red) and Cleaved Caspase-3 (Green). (B-C) 10e^4^ BxPC-3 cells were treated with 1 µg STING^WT^ or STING^R284S^ mRNA-LNP, which were pre-mixed with 10 µg/ml human APOE4 protein. At 16h post-transfection, STING^WT^ and STING^R284S^ expression was confirmed by RT-qPCR (B), and the mRNA levels of the indicated genes were measured by RT-qPCR and normalized to GAPDH mRNA levels (C). The values for untreated cells (Mock) were set to 1. (D) 0.5×10e^4^ BxPC-3 cells were treated with 1 µg STING^WT^ or STING^R284S^ mRNA-LNP, which were pre-mixed with 10 µg/ml APOE4 protein. At 16h post-transfection, cell viability was measured by Titer-GLO 3D cell viability assay. (E) 10e^4^ CD8^+^T cells were treated with 1 µg STING^WT^ or STING^R284S^ mRNA-LNP, which were pre-mixed with 10 µg/ml APOE4 protein. At 16h post-transfection, cell viability was measured by the Titer-GLO 3D cell viability assay. Error bars represent SEM of three independent experiments. (ns: not significant, * P < 0.05, **P < 0.01, ***P < 0.001).

**Figure S4.**
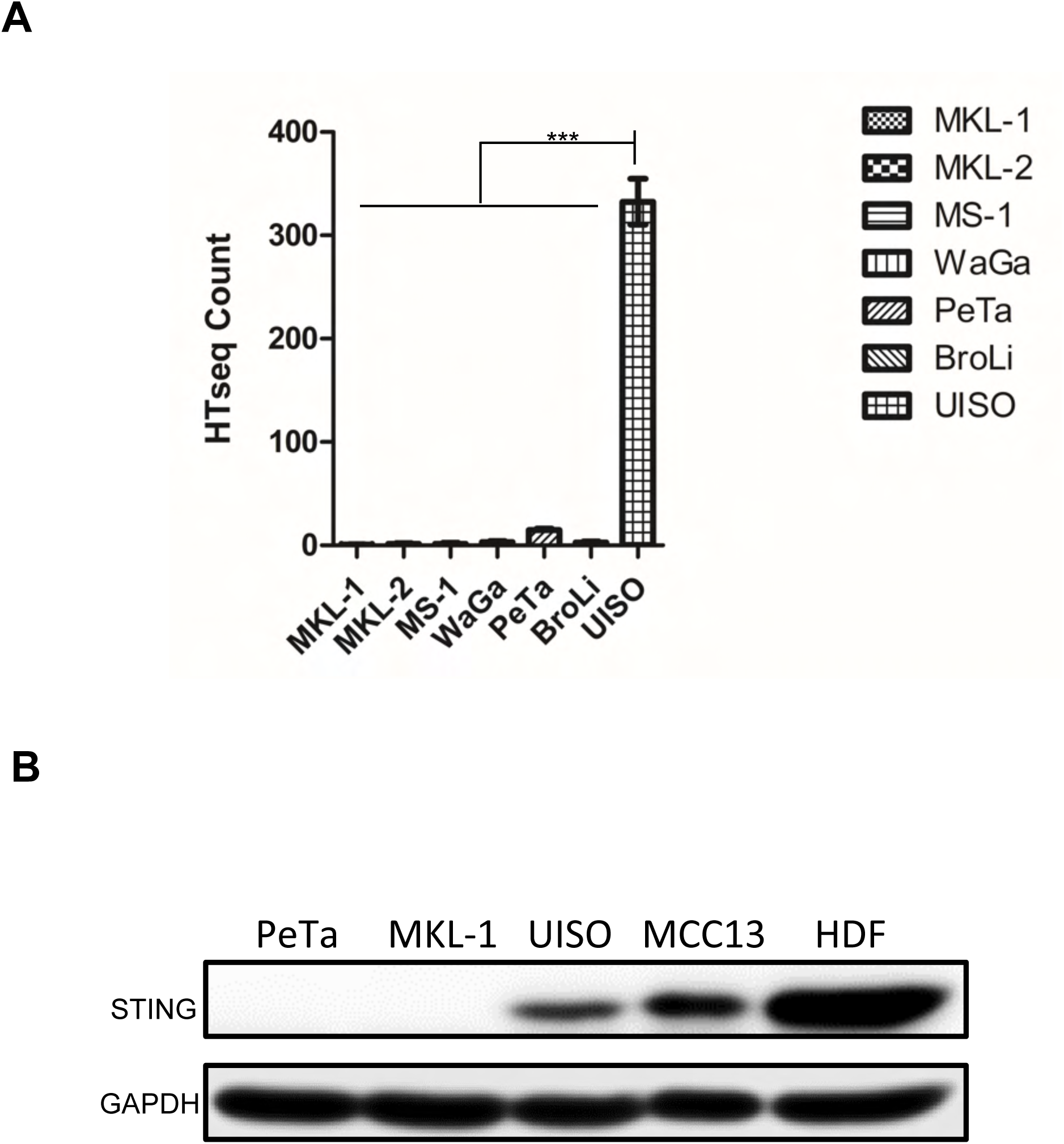
STING is silenced in classic MCPyV^+^ MCC cell lines. (A) The mRNA level (HTseq count) of STING in MKL-1 MKL-2, MS-1, WaGa, PeTa, BroLi, and UISO cells was calculated based on published RNA-seq data [87]. Error bars represent SEM of three independent experiments. (ns: not significant, * P < 0.05, **P < 0.01, ***P < 0.001). (B) Whole-cell lysates of PeTa, MS-1, UISO, MCC13 and primary HDF cells were immunoblotted using the indicated antibodies. GAPDH was used as a loading control.

**Figure S5.**
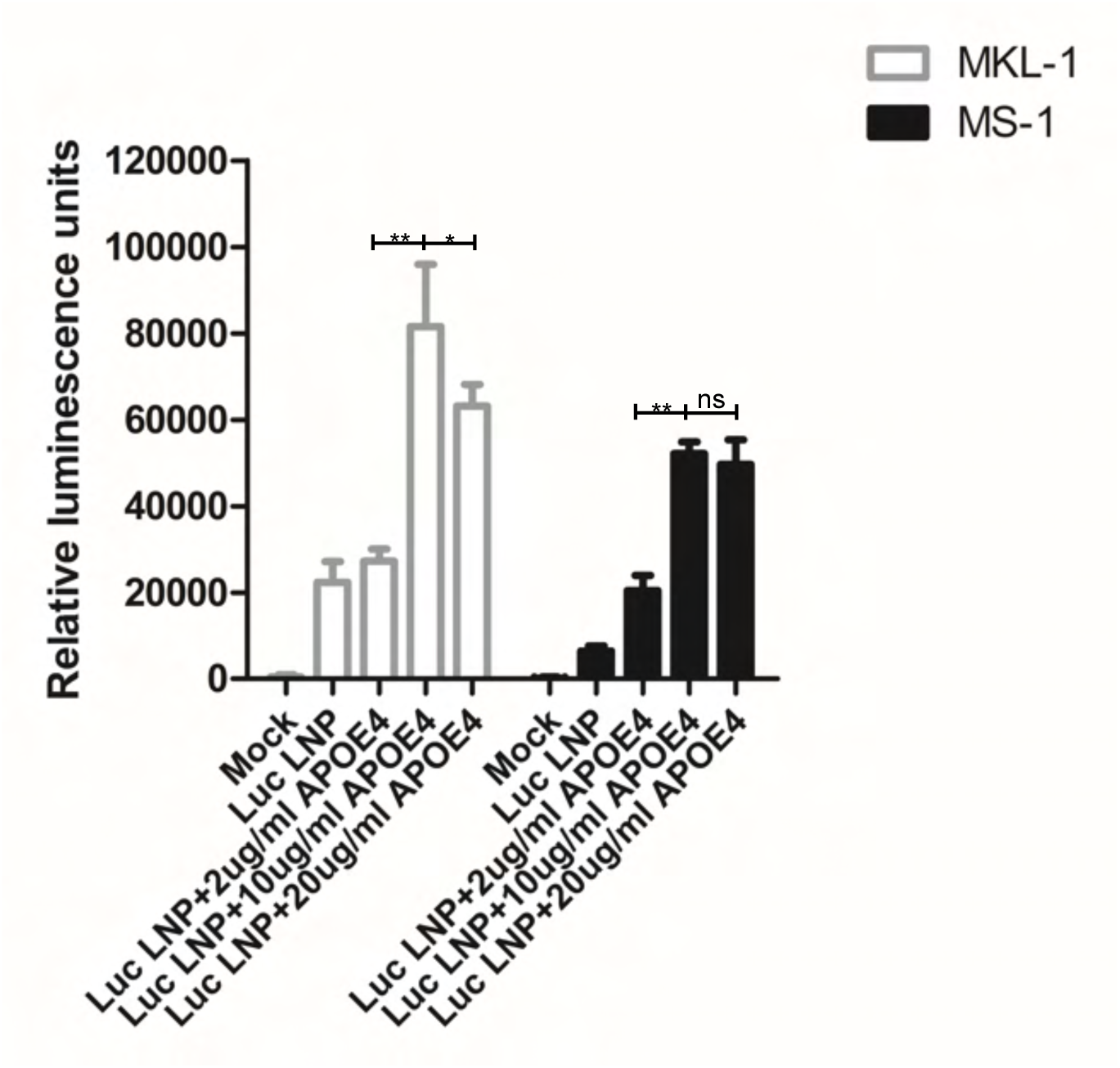
Optimizing the conditions for mRNA-LNP delivery in MCC cells. 10e^4^ MKL-1 and MS-1 cells were treated with 1 µg firefly luciferase mRNA-LNP, which were pre-mixed with the indicated concentration of the recombinant human APOE4 protein. At 16 h post-transfection, firefly luciferase activity was measured using a luciferase reporter assay system kit (Promega). (ns: not significant, * P < 0.05, **P < 0.01, ***P < 0.001).

**Figure S6.**
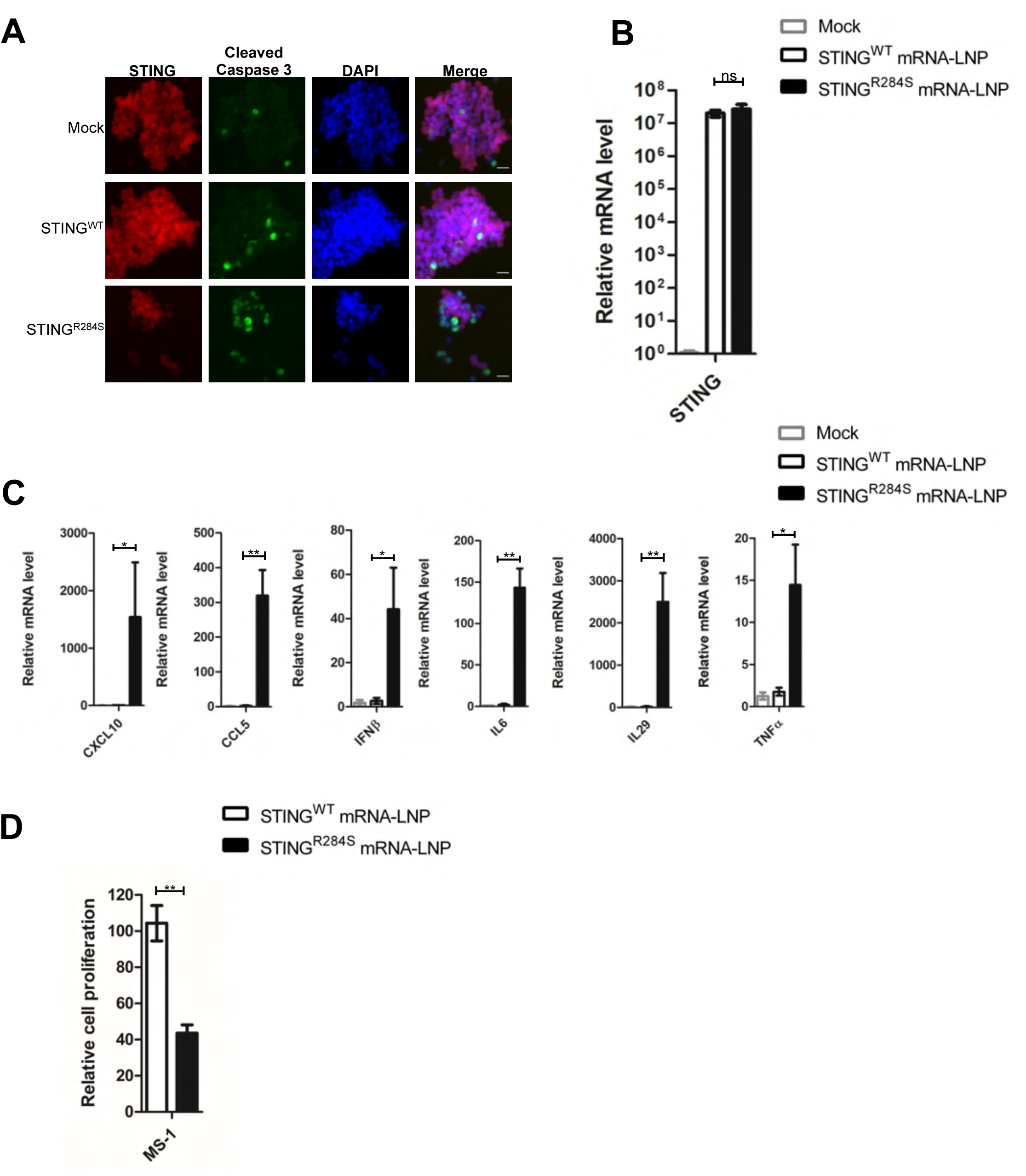
STING^R284S^ mRNA-LNP can trigger vital anti-tumor cytokine production and cell death in MCC cells. (A) 10e^4^ MS-1 cells were treated with 1 µg STING^WT^ or STING^R284S^ mRNA-LNP, which were pre-mixed with the indicated concentration of the recombinant human APOE4 protein. At 16h post-transfection, cells were stained for STING (Red) and Cleaved Caspase-3 (Green). (B-C) 10e^4^ MS-1 cells were treated with 1 µg STING^WT^ or STING^R284S^ mRNA-LNP, which were pre-mixed with 10 µg/ml human APOE4 protein. At 16h post-transfection, STING^WT^ and STING^R284S^ expression was confirmed by RT-qPCR (B), and the mRNA levels of the indicated genes were measured by RT-qPCR and normalized to GAPDH mRNA levels (C). The values for untreated cells (Mock) were set to 1. (D) 0.5×10e^4^ MS-1 cells were treated with 1 µg STING^WT^ or STING^R284S^ mRNA-LNP, which were pre-mixed with 10 µg/ml APOE4 protein at 0h and 24h. At 40h post-transfection, cell viability was measured by the Titer-GLO 3D cell viability assay. Error bars represent SEM of three independent experiments. (ns: not significant, * P < 0.05, **P < 0.01, ***P < 0.001).

